# The effect of ischaemic postconditioning on mucosal integrity and function in equine jejunal ischaemia

**DOI:** 10.1101/2020.11.25.397752

**Authors:** Nicole Verhaar, Gerhard Breves, Marion Hewicker-Trautwein, Christiane Pfarrer, Karl Rohn, Marion Burmester, Nadine Schnepel, Stephan Neudeck, Lara Twele, Sabine Kaestner

## Abstract

**Background:** Ischaemic postconditioning (IPoC) has been shown to ameliorate ischaemia reperfusion injury in different species and tissues.

**Objectives:** To assess the feasibility of IPoC in equine small intestinal ischaemia and to assess its effect on histomorphology, electrophysiology and paracellular permeability.

**Study design:** randomized controlled terminal in vivo experiment

**Methods:** Experimental jejunal ischaemia was induced for 90 min in horses under general anaesthesia. In the control group (C; n=7), the jejunum was reperfused without further intervention. In the postconditioning group (P; n=7), IPoC was implemented by clamping the mesenterial vessels after ischaemia. This was followed by 120 minutes of reperfusion in both groups. Intestinal microperfusion and oxygenation was measured during IPoC using spectrophotometry and Doppler fluxmetry. Histomorphology and histomorphometry of the intestinal mucosa were assessed. Furthermore, electrophysiological variables and unidirectional fluxrates of ^3^H-mannitol were determined in Ussing chambers. Western Blot analysis was performed to determine the tight junction protein levels of Claudin-1, Claudin-2 and Occludin in the intestinal mucosa. Comparisons between the groups and time points were performed using a two-way repeated measures ANOVA or non-parametric statistical tests for the ordinal and not normally distributed data (significance p<0.05).

**Results:** Postconditioning significantly reduced intestinal microperfusion during all clamping cycles, yet affected tissue oxygenation only during the first cycle. After reperfusion, group IPoC showed significantly less mucosal villus denudation (mean difference 21.5 %, p=0.02) and decreased mucosal-to-serosal fluxrates (mean difference 15.2 nM/cm^2^/h, p=0.007) compared to group C. There were no significant differences between the groups for the other tested variables.

**Main limitations:** small sample size, long term effects were not investigated.

**Conclusions:** Following ischaemic postconditioning, the intestinal mucosa demonstrated significantly less villus denudation and paracellular permeability compared to the untreated control group, possibly indicating a protective effect of IPoC on ischaemia reperfusion injury.

## Introduction

Small intestinal strangulation with concurrent ischemia represents a critical condition in the equine population [1]. Although many of these lesions can be treated successfully by resection and anastomosis, there is still need for alternative strategies to ameliorate ischaemia reperfusion injury in intestinal segments that are still vital and not to be resected.

The concept of ischaemic postconditioning (IPoC) describes the re-occlusion of blood supply in multiple cycles directly after an ischaemic event [2]. It has been shown to ameliorate ischaemia reperfusion injury in different species and tissues [2-5]. In human medicine, this is mainly implemented in the fields of cardiology or neurology for acute myocardial or cerebral infarction [2; 5; 6]. Most research on intestinal IPoC has been performed in rat models, demonstrating a variety of positive effects like reduced histomorphological injury, reduced apoptosis and less tissue edema [4; 7-10]. Several mechanisms have been suggested as mode of action, such as the upregulation of Hypoxia Inducible Factor-1-alpha or slower washout of protective factors as for example adenosine and aldose reductase [4; 7; 11].

The re-occlusion of blood supply after the resolution of small intestinal strangulating lesions may represent a feasible therapeutic strategy in equine colic surgery. To the authors’ knowledge, there are no reports of IPoC in intestinal ischaemia in horses. Therefore, the objective of the study was to assess the feasibility of IPoC in equine jejunum, and to evaluate the effect of this treatment strategy on different variables of ischaemia reperfusion injury in experimental equine jejunal ischaemia. The authors hypothesized that IPoC clamping effectively reduces intestinal microperfusion in the equine jejunum, and that IPoC would ameliorate ischaemia reperfusion injury.

## Materials and methods

### Animals

The study was reviewed and approved by the Ethics Committee for Animal Experiments of Lower Saxony, Germany, according to the German Animal Welfare Act. In this terminal in vivo experimental trial, 16 horses were randomly assigned to a group undergoing ischemic postconditioning (group IPoC), and an untreated control group (group C). Two horses were excluded from the data analysis for later mentioned reasons, resulting in seven horses per group. Group C consisted of five Warmbloods, one Thoroughbred and one Islandic horse. The mean age was 12.6 ± 8.7 years and the mean weight 535 ± 89 kg. Group IPoC consisted of four Warmbloods, one Thoroughbred, one Standardbred and one Islandic pony with a mean age and weight of 10.4 ± 8.6 years and 506 ± 96 kg. All horses were systemically healthy, and had been elected for euthanasia due to problems unrelated to the gastrointestinal tract.

### Anaesthesia

Before surgery, feed was withheld for 6 hours. The horses were premedicated with 0.7 mg/kg BW xylazine (Xylavet 20 mg/ml^a^), and general anaesthesia was induced with 0.1 mg/kg BW diazepam (Ziapam 5mg/kg^b^) and 2.2 mg/kg ketamine (Narketan^c^). Anaesthesia was maintained with isoflurane (Isofluran CP^a^) in 100% oxygen, and continuous monitoring of cardiovascular and respiratory parameters was performed to ensure adequate oxygenation and perfusion during the experiment. Continuous rate infusion with lactated Ringer’s solution (Ringer-Laktat EcobagClick^d^) and dobutamine (Dobutamin-ratiopharm 250mg^e^) were given to effect, to maintain the mean arterial blood pressure between 60 and 80 mmHg. One horse experienced severe anaesthetic problems unrelated to the surgical procedure, leading to exclusion from the study.

### Surgical procedure

After induction of anaesthesia, the horses were positioned in dorsal recumbency, and a routine pre-umbilical ventral midline laparotomy was performed. Thirty minutes after induction, ischemia was induced in a 1 meter jejunal segment by separately occluding both the intestinal loops and the mesentery with associated vessels with umbilical tape. During this procedure, the intestinal microperfusion was monitored by microlightguide spectrophotometry and Doppler fluxmetry (O2C^f^)[12], and the ligature was tied when the intestinal blood flow was reduced to 10 percent of the baseline, creating a 90% low-flow ischaemia. After 90 minutes of ischemia, the ligature was cut and in group C the intestines were reperfused without further intervention. In group IPoC, a delayed reperfusion was performed through re-occlusion of the mesenterial vessels by clamping for 3 cycles of 30 seconds, each followed by 30 seconds of reperfusion. For this purpose, two large haemostatic forceps were used, the jaws covered with Foley catheters to prevent trauma to the vessels (Fig. 1). This technique was established in the first horse undergoing IPoC, leading to some small alterations in clamping technique. To maintain a uniformly postconditioned test group, it was decided to exclude this horse from analysis. During the clamping cycles, the intestinal microperfusion and oxygenation were measured continuously at 0.5 Hz. The last measurement of each clamping or reperfusion cycle was designated to determine the end effect of that cycle. After the last clamping cycle, the intestines were reperfused for 120 minutes. Subsequently, the horses were euthanized without regaining consciousness, and transferred to the institute of anatomy for educational purposes.

**Figure 1:**
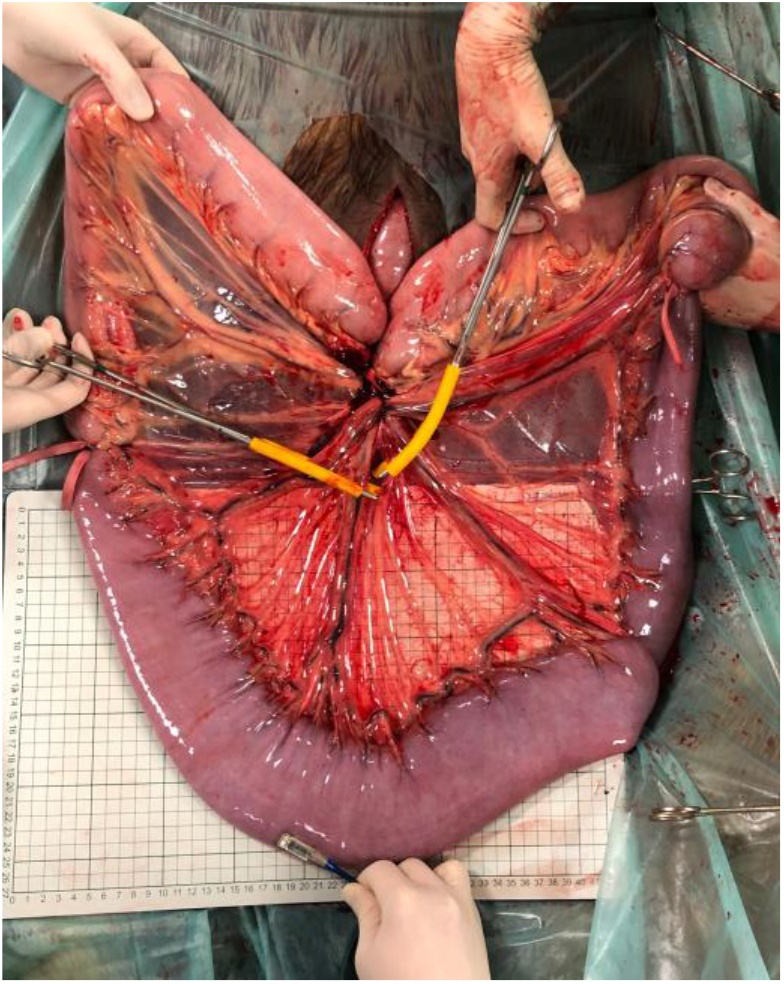
Photograph of the experimental situs showing the segmental jejunal ischaemia, with clamps positioned for postconditioning. At the antimesenterial side of the intestine, the O2C probe is in place to measure the oxygenation and microperfusion of the intestinal tissue.

### Sample collection

Full thickness intestinal segments were taken just before ischaemia (pre-ischemia sample P), at the end of ischaemia (ischaemia sample I), and at the end of reperfusion (reperfusion sample R). A second intestinal segment was taken just oral from the previously occluded ischaemic area (pre-stenotic sample PR) as a second control sample. In group IPoC, a mesenterial tissue sample was taken from the area where the clamping was executed. In group C, this sample was taken from the corresponding location in the mesentery.

### Histology and immunohistochemistry

One section of the intestinal sample was fixed in formalin, routinely processed for histopathological examination, and stained with haematoxylin and eosin (H&E). All slides were scanned to a digital format (Axio Scan.Z1^g^), and subsequently evaluated using the accompanying software (Zen 3.0 Blue edition^g^). One section per sample was evaluated by one observer who was blinded for the identity of the slides. Ten villi that were sectioned in proper alignment with their length-axis, were evaluated for histomorpology and histomorphometry. Each selected villus was scored for epithelial separation (EPS) and haemorrhage (HS) using a modified Chiu score (Table 1)[9; 13; 14]. The following histomorphometrical measurements were performed: villus and crypt height, epithelial-covered villus height, and maximal villus width. From these values the following variables were calculated: villus-to-crypt ratio, relative villus height compared to pre-ischaemia (villus height/pre-ischemic villus height), relative villus width compared to pre-ischaemia (villus width/pre-ischaemic villus width), and the percentage of denuded villus surface area as described previously [15]. The EPS and histomorphometrical measurements of each villus were averaged per slide. The HS was scored separately for the complete section. The mesentery samples were stained with H&E and evaluated for inflammation and injury of the mesenterial vessels.

**TABLE 1:**
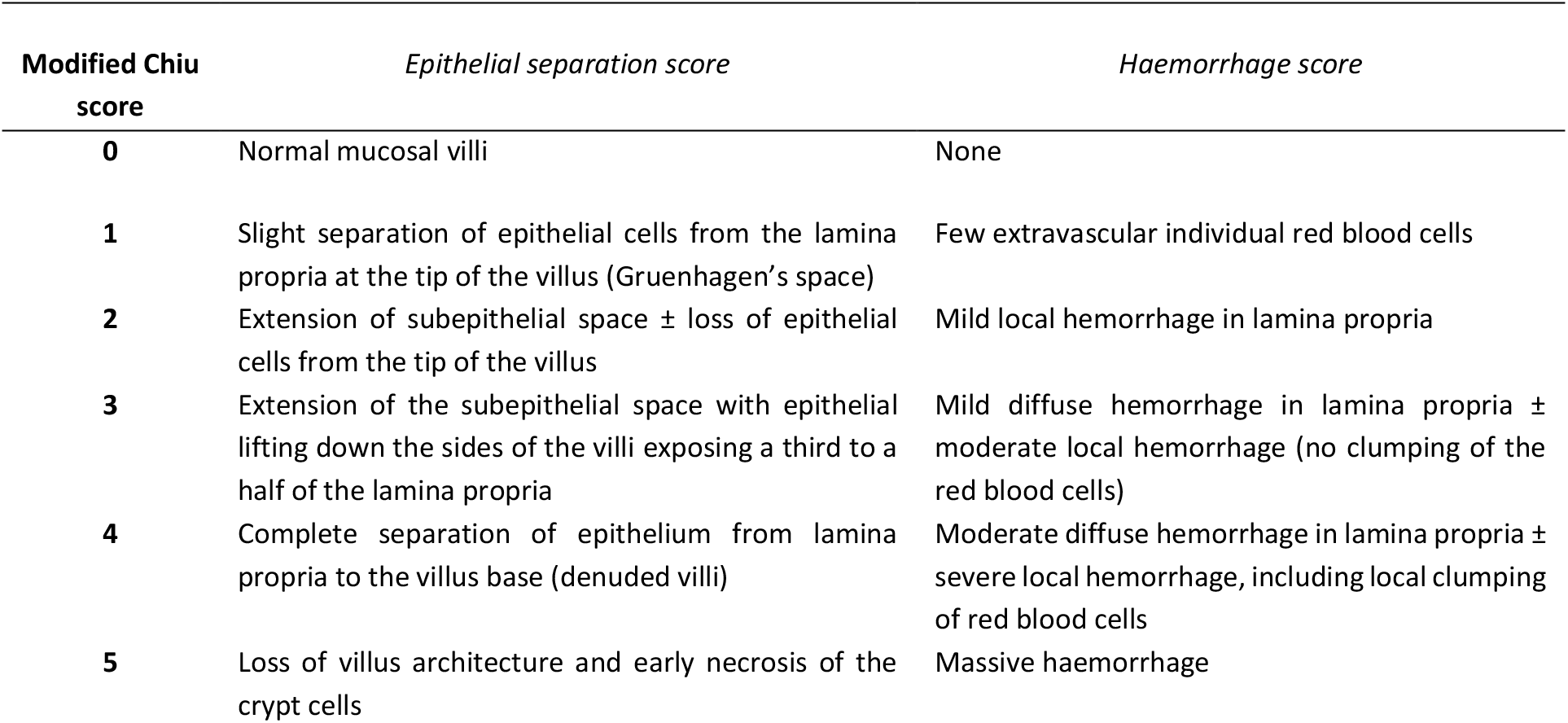
Description of the modified Chiu score assessing the histomorphology of intestinal mucosa

### Ussing chambers

The intestinal samples from 11 horses were also subjected to electrophysiological and permeability measurements. The samples of the first 4 horses were not subjected to this analysis due to the availability of the Ussing chambers during the experiment. The mucosa was stripped of the seromuscular layer and mounted in Ussing chambers with a tissue exposing area of 1.13 cm^2^, and bathed in a modified Krebs-Henseleit-buffer aerated with carbogen as previously reported [16]. In the absence of electrical and chemical gradients, short circuit currents (I_sc_ in μEq/cm/h) and transepithelial potential differences (PDt) were measured through a computer-controlled voltage clamp device^h^. Tissue conductance (Gt) was determined from the changes in PDt elicited by bipolar current pulses of 100 μA/cm^2^. Fluid resistance and junction potential were measured before the mucosa was mounted, and this was corrected for during the experimental period. Measurements were performed in the pre-ischaemia, ischaemia and reperfusion samples under basal conditions, and after the addition of 10 mM alanine or 10 mM glucose to the luminal side, to evaluate the sodium-dependent alanine and glucose transport based on the response in short circuit currents. 10^-5^M Forskolin^i^ was added at the end of the experimental period to confirm tissue viability.

In different chambers, 4 μCi of ^3^H-mannitol^j^ was added after an equilibration period of 15 min, to determine the unidirectional flux rates as a measure for paracellular permeability. Radioactivity was measured as % disintegrations/min using a liquid scintillation counter (Packard Tri-Carb liquid scintillation analyzer^i^) in 250-μL samples collected from both sides of the chamber at 15-minute intervals. Mucosal-to-serosal (J_ms_), serosal-to-mucosal (J_sm_) and netto flux rates (J_net_ = J_ms_ - J_sm_) of mannitol were calculated from tracer appearance using standard equations.

### Western blot analysis for tight junction protein

Intestinal mucosa samples were snap frozen in liquid nitrogen and stored at −80°C until further processing. The frozen tissue was homogenised with zirconia balls using a high-speed homogenizer (FastPrep-24™ 5G^k^) in a lysis buffer containing 250mmol/l sucrose, 20mmol/l TRIS, 5mmol/l EGTA and 5mmol/l MgSO_4_*7H_2_O at pH 7.5 with freshly added protease inhibitor cocktail tablets (cOmplete ULTRA Tablets^l^) This step was followed by two centrifugations (2000g, 20min at 4°C and 40000g, 60min at 4°C). After discarding the supernatant, the pellet was resuspended with 10mmol/l TRIS-buffer pH 7.4 containing 150mmol/l NaCl as well as protease inhibitor cocktail tablets. Protein concentrations were measured with a commercial protein assay using Bradford reagent^m^.

Two μg of intestinal crude membranes for Claudin-1 and 10 μg for Claudin-2 were separated by 12% SDS-Page. For Occludin 2μg intestinal crude membranes were separated by 8.5% SDS-Page. Proteins were transferred to nitrocellulose membranes^n^. Membranes were blocked in Tris-Buffered Saline containing 0.1% Tween 20 (TBS/T) (Claudin-1) or Phosphate-Buffered Saline containing 0.1% Tween 20 (PBS/T) (Claudin-2 and Occludin) with 5% fat-free milk powder. Subsequently, the membranes were incubated overnight at 4°C with the primary antibody in TBS/T with 5% fat-free milk powder (anti-Claudin-1°), PBS/T with 3% bovine serum albumin (anti-Claudin-2^p^) or PBS/T (anti-Occludin^q^). After washing, the membranes were incubated with secondary anti-rabbit antibody^i^ 5% fat-free milk powder in TBS/T (Claudin-1) or PBS/T (Occludin), or with secondary anti-mouse horseradish peroxidase-conjugated antibody^i^ in PBS/T with 2% fat-free milk powder (Claudin-2). Membranes that were incubated with the secondary antibody only, served as a negative control to ensure that no nonspecific signals were detected. The proteins were detected with a chemiluminescence detection and imaging system (ChemiDoc^r^), and the densitometric measurements were performed using the accompanying software. For semi-quantification of the proteins, the amount of the investigated proteins was normalized to the amount of total protein per lane. Sample analysis was performed in duplicates, and the mean of both assays was used for further data analysis. The data were expressed as percentage compared to pre-ischaemia.

### Data analysis

A power analysis was performed prior to commencing the study using free available software (G*Power 3.1.9.1^s^). To detect a difference of 0.5 grade in the histomorphology score between the treatment groups with a standard deviation of 0.3, based on a power of 0.8 and alpha of 0.05, a total sample size of 14 horses was required.

Statistical analysis and graph design were performed with commercial software (SAS 9.4m5 with the Enterprise Guide Client 7.15^t^, and Graphpad Prism 8.3.1^u^). The data were tested for normal distribution by visual assessment of the qq-plots of the model residuals and the Shapiro-Wilks-test was performed. Variance homogeneity was investigated by visual assessment of the homoscedasticity plots and by performing Levene’s test. The normally distributed variables were expressed as mean (± standard deviation), and the non-parametric as median (min-max). P-values of <0.05 were considered significant. The ordinal variables (histomorphology score) and not normally distributed data (intestinal microperfusion and saturation) were analysed using distribution free models for independent (treatment and control group) and correlated effects (time points). A Mann-Whitney-U Test was used to compare the results between the different groups at each time point. For comparing the correlated different time points, a permutation test (as exact Friedman test) for repeated measures was used [17], with a post-hoc Sidak-test for multiple pairwise comparisons.

For analysis of the normally distributed data (histomorphometry, electrophysiology, flux rates, and western blot results), a two-way analysis of variance (ANOVA) was performed for one independent effect (group), and the time points as repeated effect. This was implemented to compare the values between the different time points and groups, with the horses as subject effect. The p-values were subjected to the Greenhouse-Geisser correction. Post-hoc multiple pairwise comparisons were performed between the groups with Sidak’s test, and between the time-points within one group with Tukey’s test.

## Results

### Intestinal microperfusion and oxygenation

Pre-ischaemia, the median intestinal microperfusion of all horses was 353 (209 – 488) Arbitrary Units (AU), and was significantly reduced to 39 (21 – 66) AU after tightening the ligature (p<0.0001). At the end of ischaemia, this was 47 (32 – 60) AU. Tissue saturation was 92.5 (75.7 – 98.3) % at pre-ischaemia, and reduced to 56.7 (12.1 – 75.6) % at the beginning of ischaemia, and was 25.0 (14.2 – 44.3) % at the end of ischaemia. The microperfusion and saturation elicited by the experimental ischaemia did not differ between the groups.

In group IPoC, postconditioning significantly reduced the blood flow during all clamping cycles to an average of 34.5 AU (p<0.001)(Fig. 2). A significant decrease in saturation only occurred during the first clamping cycle with a median difference of 47.1 % (p=0.02), which was also significantly lower than the saturation during the third cycle (p=0.01). During the 30 sec reperfusion cycles, microperfusion and oxygenation returned to the level of the pre-ischaemia measurement.

**Figure 2:**
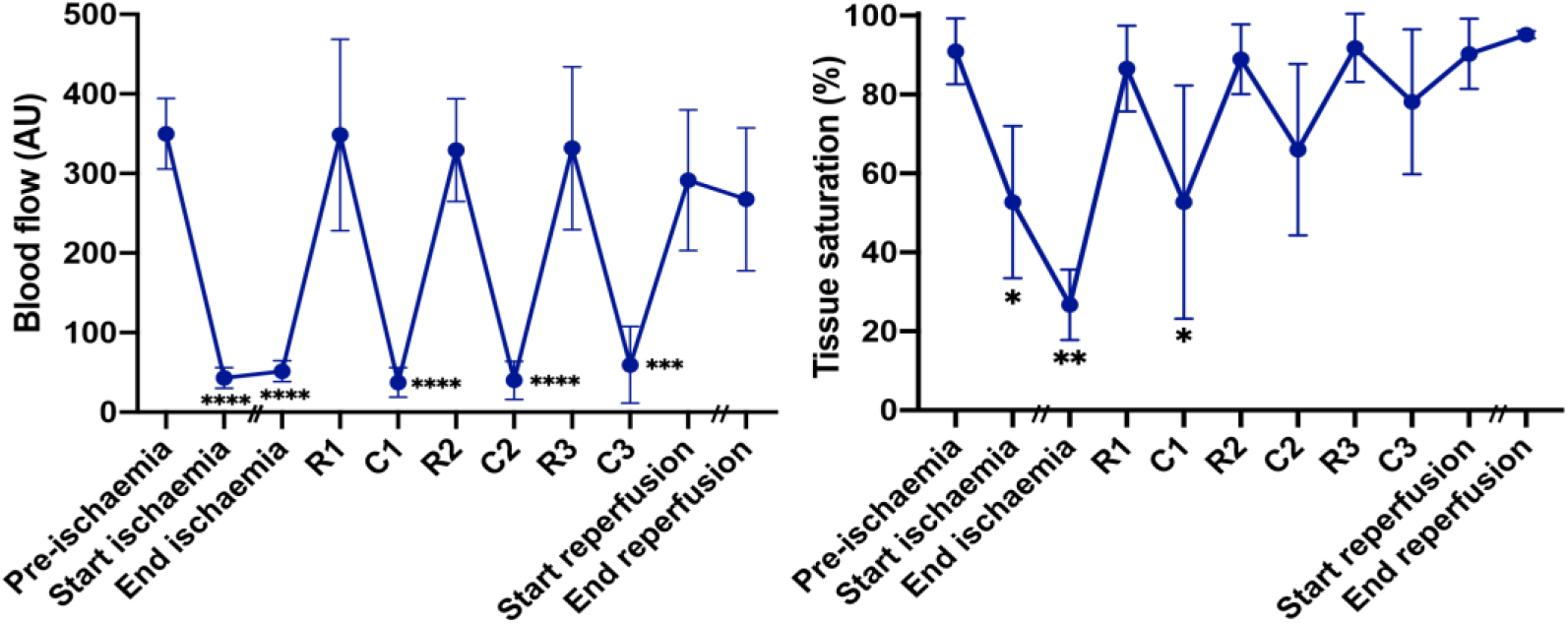
Diagram showing (A) intestinal microperfusion in Arbitrary Units and (B) oxygenation in % measured in the equine jejunum of horses undergoing postconditioning with three clamping (C) and release (R) cycles of 30 seconds each. Data are presented as median (min-max), time points that differ from the pre-ischaemic measurement are marked with an asterisk *(* = p<0.05, ** = p<0.01, *** = p<0.001, **** = p<0.0001)*.

### Histomorphology score

In the pre-ischaemia samples, all horses showed an EPS of 0 and a HS of 0 or 1 (Fig. 3). There was a significant increase of both scores during ischaemia (EPS p= <0.0001 and 0.004, HS p= 0.001 and 0.007 for group C and IPoC, respectively), and no change occurred during reperfusion. There were no significant differences between the groups during ischaemia (p=0.10) or reperfusion (p=0.07) in EPS, and the same was found for the HS (p=0.64 and 0.59, respectively). The pre-stenotic sample taken at the end of the experiment showed an EPS of 0 in all horses, and a HS of 0 in all horses except for one horse in each group with a score of 1.

**Figure 3:**
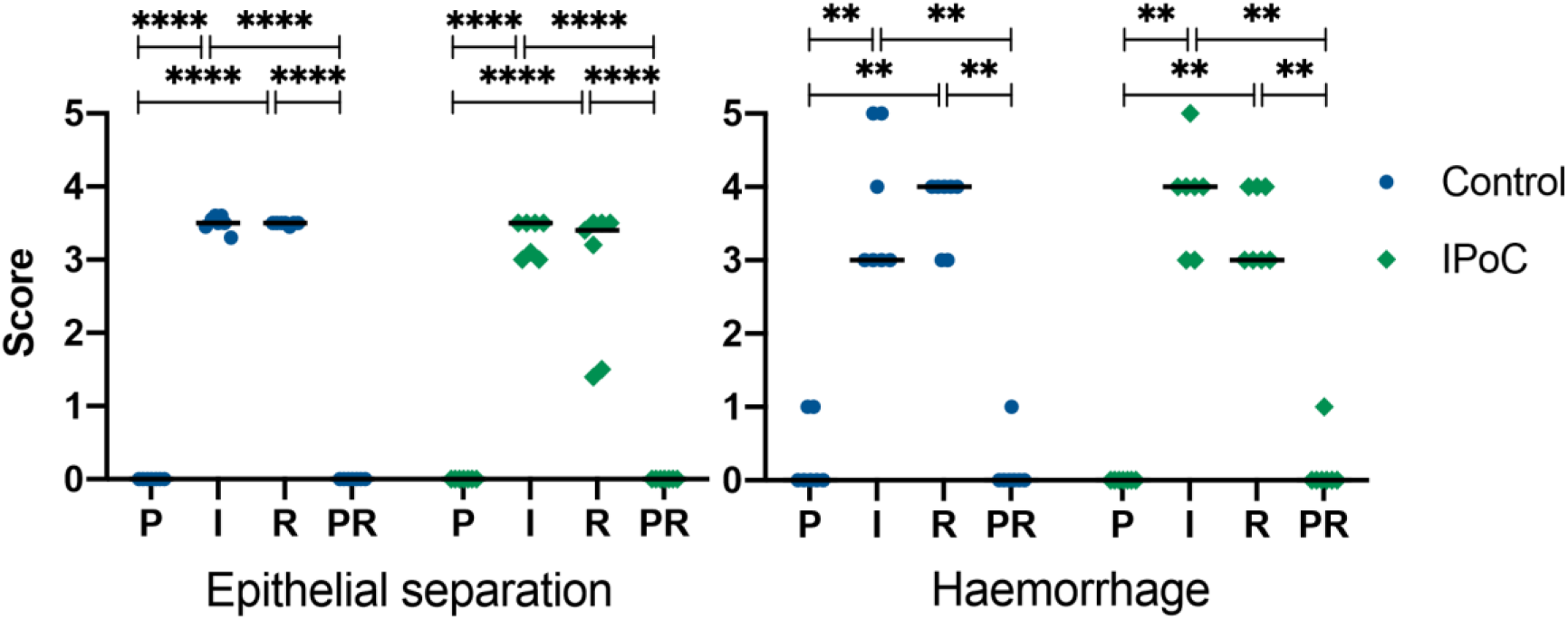
Scatterplot diagram of a modified Chiu score for mucosal histomorphology during pre-ischaemia (P), ischaemia (I), reperfusion (R), and in a prestenotic control sample (PR) in an experimental model of equine jejunal ischaemia. C = control group; IPoC = group undergoing ischaemic postconditioning. The horizontal bar displays the median. Significant differences are marked with an asterisk *(* = p<0.05, ** = p<0.01, *** = p<0.001, **** = p<0.0001)*.

### Histomorphometrical measurements

The percentage of denuded villus surface area was 0% in all P and PR samples (Table 2). After ischaemia, this was 57 ± 15% in group C and 39 ± 5% in group IPoC (p = 0.09). After reperfusion, group C showed significantly more villus denudation compared to group IPoC (mean difference 21.5 %, CI 2.64 – 40.4, p = 0.02). The villus-crypt ratio and the relative villus height and width did not show significant differences between the treatment groups in any of the time points. These variables were all significantly affected by ischaemia, without further progression during reperfusion (Table 2).

**TABLE 2:** Histomorphometry results

|  | Pre-ischæmia |  | Ischæmia |  | Reperfusion |  | Prestenotic |  |
| --- | --- | --- | --- | --- | --- | --- | --- | --- |
|  | C | IPoC | C | IPoC | C | IPoC | C | IPoC |
| Denuded villus surface area (%) | 0 | 0 | 57 ± 15<sup>b</sup> | 39 ± 5<sup>b</sup> | 49 ± 11<sup>a,b</sup> | 28 ± 13<sup>a,b</sup> | 0 | 0 |
| Villus-crypt ratio | 2.1 ± 0.3 | 2.1 ± 0.3 | 1 ± 0.1<sup>b</sup> | 1.3 ± 0.3<sup>b</sup> | 0.9 ± 0.1<sup>b</sup> | 1.2 ± 0.3<sup>b</sup> | 2.0 ± 0.3 | 2.1 ± 0.3<sup>b</sup> |
| Relative villus height (%) | 100 | 100 | 37 ± 9<sup>b</sup> | 54 ± 16<sup>b</sup> | 29 ± 4<sup>b</sup> | 44 ± 21<sup>b</sup> | 89 ± 11 | 80 ± 10<sup>b</sup> |
| Relative villus width (%) | 100 | 100 | 171 ± 19<sup>b</sup> | 142 ± 24<sup>b</sup> | 153 ± 22<sup>b</sup> | 159 ± 38<sup>b</sup> | 113 ± 15 | 114 ± 7 |

Data are expressed as mean ± standard deviation. A significant difference between the treatment groups is marked with a matching uppercase letter ‘a’, and significant differences of a time point compared to the pre-ischæmia measurement are marked with an uppercase letter ‘b’ (p<0.05). C = control group; IPoC = group undergoing ischaemic postconditioning.

### Mesentery

The mesentery of both control and postconditioned horses showed hyperaemia and mild local haemorrhage around the vessels. The vessel walls were intact, and a moderate amount of neutrophils was seen in and around the mesenterial vessels. No differences could be detected between the postconditioned mesentery and that of the control horses.

### Electrophysiology

One horse showed unusually high values for tissue conductance in all samples. Therefore, this horse, belonging to group C, was excluded from further analysis of the electrophysiology and flux rates. This analysis was conducted with the remaining 10 horses: 4 in group C and 6 in group IPoC. The values for short circuit currents and tissue conductance are summarized in supplementary item 1.

Under basal conditions, the mean I_sc_ values in the different chambers ranged between -0.4 and 0.3 μEq/cm^2^/h. In the pre-ischaemia sample, the short circuit current increased to 3.4 ± 0.8 μEq/cm^2^/h in group C and 3.1 ± 1.1 μEq/cm^2^/h in group IPoC in response to the addition of alanine, and to 2.5 ± 0.6 μEq/cm^2^/h and 2.3 ± 0.8 μEq/cm^2^/h after the addition of glucose (Fig. 4). Compared to this time point, the ischaemia samples showed significantly less response to the addition of alanine (group C: mean difference 3.2, CI 1.26 to 4.8, p=0.008; group IPoC: mean difference 2.4, CI 1.2 to 3.7, p=0.003) and glucose (group C: mean difference 2.4 μEq/cm^2^/h, CI 1.2 to 3.6, p=0.007; group IPoC: mean difference 1.8, CI 1.0 to 2.6, p=0.002). After reperfusion, the short circuit current response to alanine was comparable to ischaemia, and there were no differences between the treatment groups in any of the chambers or time points.

**Figure 4:**
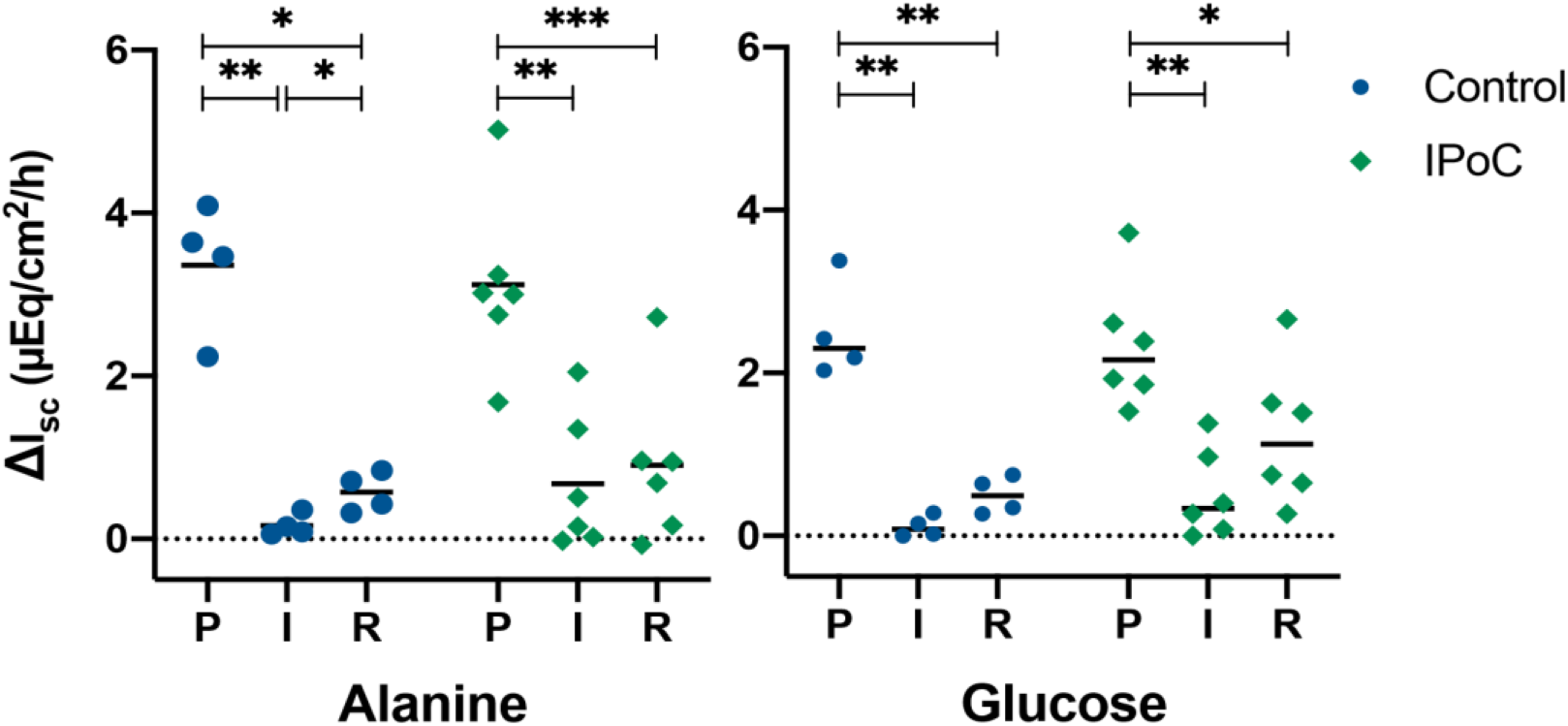
Scatterplot diagram displaying the change in short circuit currents (I_sc_) in μEq/cm^2^/h measured in Ussing chambers after the addition of alanine or glucose during pre-ischaemia (P), ischaemia (I) and reperfusion (R) in an experimental model of equine jejunal ischaemia. C = control group; IPoC = group undergoing ischaemic postconditioning. The horizontal bar displays the mean. Significant differences are marked with an asterisk *(* = p<0.05, ** = p<0.01, *** = p<0.001, **** = p<0.0001)*.

During pre-ischaemia, the tissue conductance ranged between 13.7 – 26.1 mS/cm^2^ in the alanine and glucose chambers under basal conditions. The addition of alanine, glucose or forskolin did not affect the tissue conductance, and there were no significant differences between the groups for any of the time points. After ischaemia, there was a significant increase in tissue conductance (group C: mean difference -5.9 mS/cm^2^, CI -11.3 to -0.5, p=0.04; group IPoC: mean difference -6.0 mS/cm^2^, CI -10.1 to -1.9, p=0.01) without further changes during reperfusion.

### ^3^H-mannitol flux rates

Pre-ischaemia, J_ms_ was 21 ± 7.6 and 20 ± 6.3 nM/cm^2^/h in group C and IPoC, respectively (Fig. 5). There was a significant increase in both groups during ischaemia (group C: mean difference -19.6 nM/cm^2^/h, CI -33.4 to -5.9, p = 0.02; group IPoC: mean difference -15.0 nM/cm^2^/h, CI -22.5 to -7.5, p = 0.003). Only group C showed further progression during reperfusion (mean difference -13.0 nM/cm^2^/h, CI -21.0 to -4.10, p = 0.02). Moreover, group C showed a significantly higher J_ms_ compared to group IPoC at this time point (mean difference 15.2 nM/cm^2^/h, CI 4.9 to 25.5, p = 0.007).

**Figure 5:**
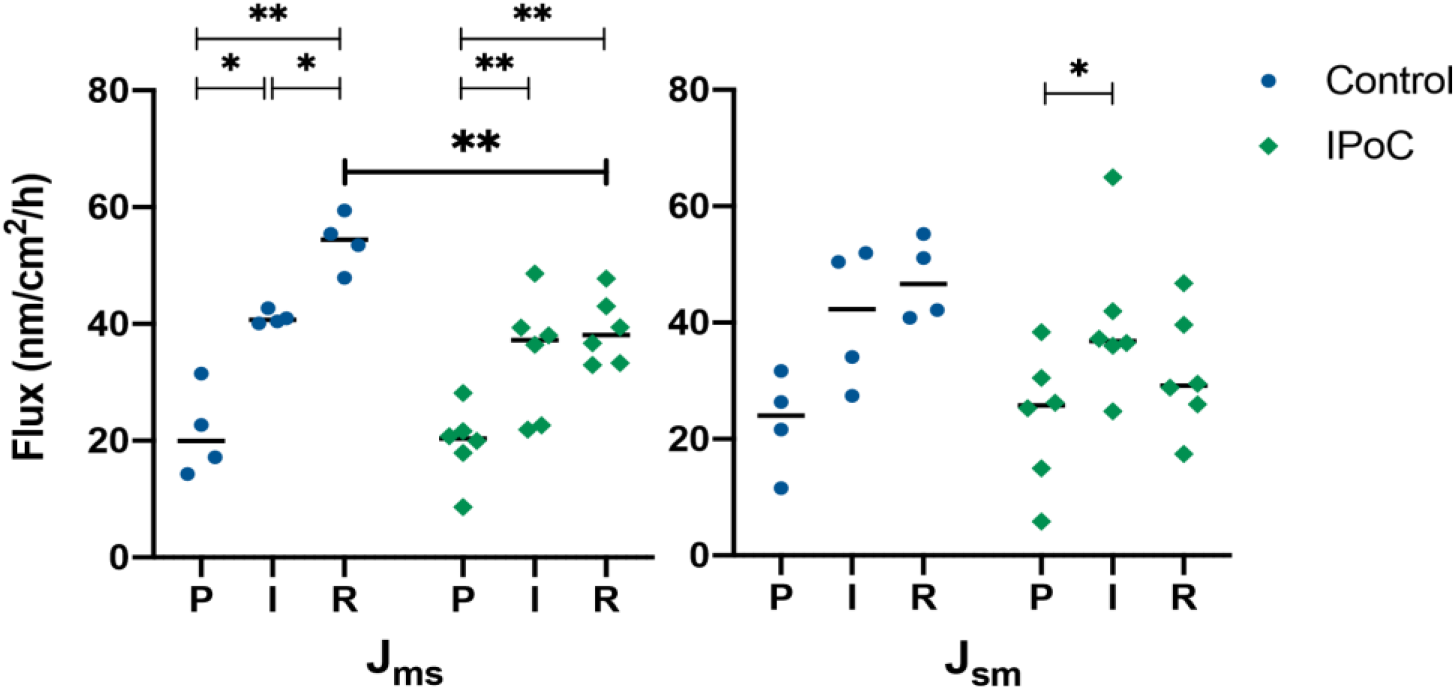
Scatterplot diagram of the unidirectional fluxrates of H^3^-mannitol measured in Ussing chambers during pre-ischaemia (P), ischaemia (I) and reperfusion (R) in an experimental model of equine jejunal ischaemia. Mucosal-to-serosal (J_ms_) and serosal-to-mucosal (J_sm_) were calculated from tracer appearance in nM/cm^2^/h. The horizontal bar displays the mean. Significant differences are marked with an asterisk *(* = p<0.05, ** = p<0.01, *** = p<0.001, **** = p<0.0001)*. C = control group; IPoC = group undergoing ischaemic postconditioning.

J_sm_ was 23 ± 8.5 nM/cm^2^/h in group C and 24 ± 11.5 in group IPoC during pre-ischaemia (Fig. 6). Group IPoC showed a significant increase during ischaemia (mean difference -16.7 nM/cm^2^/h, CI -32.5 to -0.9, p = 0.04). The reperfusion samples did not show significant differences compared to the other time points, and there were no significant differences between the groups.

**Figure 6:**
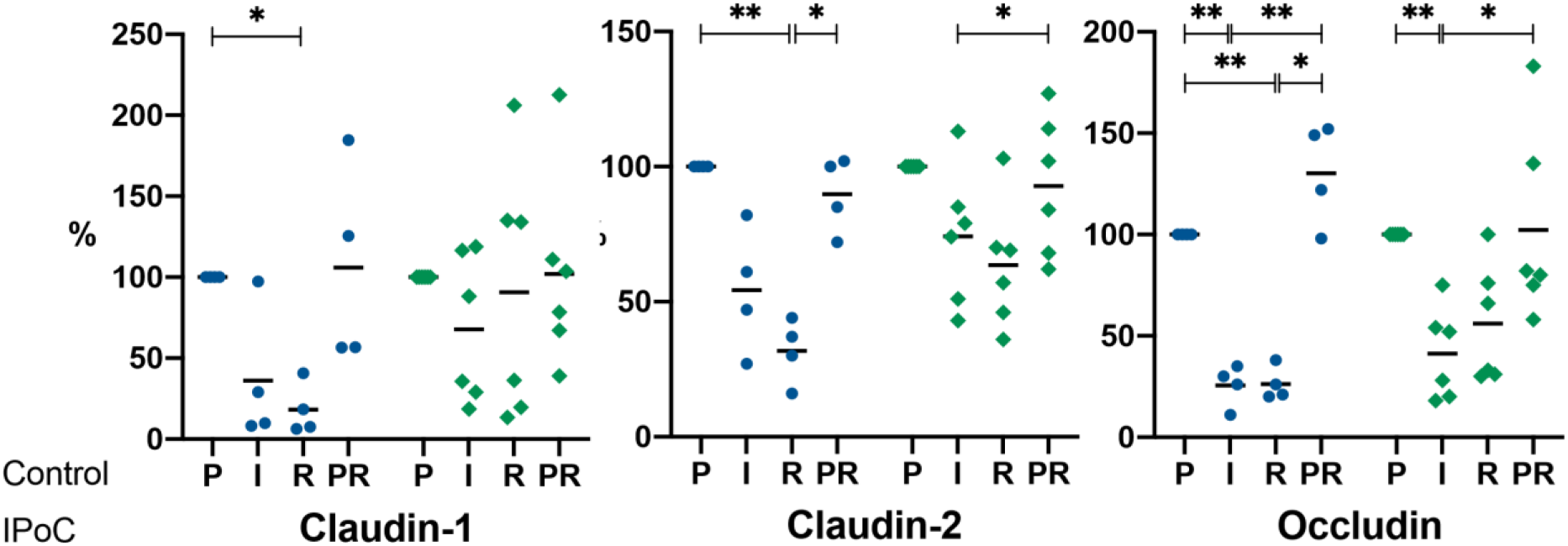
Scatterplot diagram of the protein levels of (A) Claudin-1, (B) Claudin-2, and (C) Occludin analysed by western blotting of the intestinal mucosa sampled during pre-ischaemia (P), ischaemia (I), reperfusion (R), and a prestenotic control sample (PR) in an experimental model of equine jejunal ischaemia. C = control group; IPoC = group undergoing ischaemic postconditioning. The horizontal bar displays the mean. Significant differences are marked with an asterisk *(* = p<0.05, ** = p<0.01, *** = p<0.001, **** = p<0.0001)*.

Pre-ischaemia, J_net_to was -1.4 ± 3.3 and -4.0 ± 8.3 nM/cm^2^/h in group C and IPoC, respectively. The net fluxrate did not change significantly over time, and there were no differences between the groups.

### Tight junction proteins

The Claudin-1 and -2 protein levels did not change significantly during ischaemia (Fig. 6A and B). Group C showed a significantly lower protein levels during reperfusion compared to the pre-ischaemia sample for both Claudin-1 (mean diff. 82%, CI 32 to 131, p = 0.01) and Claudin-2 (mean diff. 68%, CI 31 – 105, p = 0.009), which could not be detected in group IPoC. The Claudin-2 protein level during reperfusion was also significantly decreased compared to the prestenotic sample in group C (mean diff. -58%, CI - 98 to -18, p = 0.02). In group IPoC, the Claudin-2 protein level during ischaemia was significantly decreased compared to the prestenotic sample (mean diff. -19%, CI -34 to -3, p = 0.02).

The Occludin protein level exhibited a significant decline during ischaemia in both groups (group C: mean diff. 75%, CI 43 to 107, p = 0.004; group IPoC: mean diff. 59%, CI 20 to 98, p = 0.009). Only group C showed a significantly lower Occludin level during reperfusion compared to pre-ischaemia (mean diff. 74%, CI 48 to 99, p = 0.002). The prestenotic sample had significantly higher occludin levels compared to both ischaemia and reperfusion in group C (mean diff. -105%, CI -160 to -49, p = 0.008 and mean diff. -104%, CI -198 to -10, p = 0.04, respectively), and compared to ischaemia only in group IPoC (mean diff. -61%, CI -113 to -9, p = 0.01).

There were no significant differences for any of the proteins in the direct comparison between the groups, and the prestenotic samples did not differ from the pre-ischaemia samples.

## Discussion

This is the first study describing the implementation of ischaemic postconditioning in the horse. The main findings were that postconditioning clamping effectively reduced the intestinal microperfusion during all clamping cycles and that postconditioning resulted in a significantly lower paracellular permeability as well as less epithelial denudation after reperfusion compared to the untreated control group. There were no differences between the two groups for the other tested variables.

In the pre-ischaemia samples, basal I_sc_ and G_t_ as well as the response to alanine or glucose were comparable to previously reported values [18]. The effect of ischaemia and reperfusion was demonstrated by an increase in tissue conductance and a reduction in active transport. Postconditioning did not affect these variables. Other IPoC studies have not investigated intestinal electrophysiological parameters and flux rates, therefore precluding a direct comparison. Both unidirectional ^3^H-mannitol fluxrates increased as an effect of ischaemia, reflecting increased paracellular permeability. Further deterioration during reperfusion was found only in group C. Moreover, group IPoC demonstrated a lower mucosal-to-serosal flux compared to group C, indicating an effect of postconditioning on paracellular permeability. The disparity between the effect on flux rate and the absence of an effect on tissue conductance and short circuit current, could be attributed to the transcellular pathway including active transport not being affected by postconditioning. Paracellular permeability is principally regulated by tight junctions, consisting of 4 different types of transmembrane proteins including occludin, claudins, junctional adhesion molecules, and tricellulin. In the current study, Claudin-1 and -2 as well as Occludin levels in the intestinal mucosa were assessed. Claudin-1 is a barrier-forming tight junction important for TJ integrity [19; 20] and the most commonly expressed claudin-protein in the equine intestine [21]. The “leaky” junction claudin-2 forms pores that regulate the paracellular movement of Na^+^, Ca^2+^ and water [22; 23]. Following ischaemia, studies have reported decreased Claudin-1, -2, -4, and -7 and occludin expression in the small intestine [24; 25]. Furthermore ZO-1, occludin and Claudin-1, -3, and -5 expression and distribution have been reported to correlate with the recovery of intestinal barrier function [26; 27]. In the current study, the control group exhibited a significant decrease all tested tight junction protein levels at reperfusion compared to pre-ischaemia, which could not be detected in group IPoC. Due to the relatively short time frame of reperfusion, changes in protein level may reflect protein damage rather than decreased protein expression. Consequently, this result may indicate an effect of postconditioning on tight junction deterioration in the mucosa. Nevertheless, no significant differences could be detected in a direct comparison between the treatment groups, although this could be the consequence of the small sample size, possibly resulting in a type II statistical error. Hence the role of tight junction proteins in the equine mucosal response to postconditioning remains uncertain.

Another explanation for the reduced tissue permeability after postconditioning, could be the smaller denuded villus surface area, exposing less lamina propria to facilitate leakage. A possible explanation for this smaller denuded area could the reduced loss of epithelial cells during reperfusion. The difference in denuded villus area was not reflected by a significant difference in epithelial separation score. This may be explained by the fact that most ischaemia and reperfusion samples in this experimental model are assigned to grade 3 of the modified Chiu score, and the score might be not sensitive enough to detect smaller differences. Histomorphometrical assessment may be more suitable to detect smaller differences as seen in the current study.

Until now, nearly all studies investigating the effect of IPoC in the intestine, were performed in rats that were anaesthetized by intraperitoneal administration of ketamine and xylazin, and subjected to complete occlusion of the cranial mesenteric artery [4; 7-10; 28; 29]. In these studies, differences in Chiu score of 1 – 2 points were found between the control and treatment groups [4; 7-10], demonstrating greater changes than those found in the current study. This may be attributed to the difference in species, anaesthetic protocol or IPoC technique. Furthermore, the difference in ischaemia model could be of significance, considering that the occlusion of the cranial mesenteric artery alone is associated with a highly variable degree of injury [30].

The main limitations of the current study are the small sample size and the lack of long term effect evaluation. Furthermore, a pharmacological pre- or postconditioning effect of isoflurane cannot be ruled out [31]. This would be equally present in both groups, but it could mitigate the potential protective effect of IPoC. The elicited ischaemia injury exhibited mild variation between the individual horses, possibly representing normal biological variation or the result of minor differences in blood flow in the ischaemia model used. Therefore, it cannot be completely excluded that differences between the groups are a result of a disparity in the progression of ischaemia reperfusion injury. Nevertheless, no significant differences were found after ischaemia, and the microperfusion and oxygenation results demonstrate consistent experimental ischaemia in both groups. One horse showed very high values for tissue conductance during all time points, leading to exclusion of the electrophysiological results and fluxrates from further analysis. The histology results did not differ from the other horses; therefore, the horse was not excluded from histomorphological analysis.

The use of modified haemostatic forceps proved to be a feasible technique for postconditioning in the equine jejunum, and the mesentery was not affected negatively by the clamping. Different algorithms of occlusion cycles have been investigated for intestinal IPoC, ranging from 2 – 6 cycles of 10 to 120 seconds each [9; 28; 29]. It was found that shorter and more frequent cycles have the greatest effect. For the current study, an algorithm of 3 cycles of 30 seconds was elected, because this is most commonly reported in the literature [4; 7; 8; 10; 32]. Up to date, oxygenation and microperfusion have not been documented during intestinal postconditioning. This study revealed that only the first clamping cycle led to significant tissue re-desaturation in this algorithm. This is most likely explained by the short time span of flow reduction, combined with a swift recovery of oxygenation during the short bouts of reperfusion. This suggests that the clamping cycle should be prolonged for a consistent reduction in oxygenation, although this may vary between species, taking differences in physiological heart rates and cardiac index into account. On the other hand, the exact mode of action of IPoC still remains unclear, and desaturation with concurrent hypoxia may not be the key mediator for the induction of its protective mechanisms [11].

In equine medicine, postconditioning could be implemented during colic surgery. This would not apply to intestine that is irreversibly damaged during ischaemia, but might be indicated in cases with mild to moderate ischaemic injury, where IPoC could potentially ameliorate further injury and decrease post-operative complications. From a practical perspective, clamping with haemostatic forceps may not be feasible in longer intestinal segments. An alternative would be the manual reocclusion of the mesenterial vessels after release of ischaemia, or conducting the initial release of ischaemia in a decelerated manner.

In conclusion, it is possible to perform postconditioning in the equine jejunum by use of haemostatic forceps, although the duration of the clamping cycle would need to be increased to establish significant desaturation in all cycles. Despite the lack of major differences in the histopathology score, the mucosa of horses undergoing IPoC demonstrated significantly lower paracellular permeability and denuded villus surface area after reperfusion compared to the untreated control group, indicating a protective effect of IPoC on ischaemia reperfusion injury. More research is needed to determine the long-term effects and the underlying protective mechanisms of ischaemic postconditioning in the equine small intestine.

## Supporting information

supplementary item 1

## Manufacturer’s details

a. CP-Pharma GmbH, Burgdorf, Germany
b. Ecuphar GmbH, Greifswald, Germany
c. Vétoquinol GmbH, Ismaning, Germany
d. B. Braun Melsungen AG, Melsungen, Germany
e. Ratiopharm GmbH, Ulm, Germany
f. LEA Medizintechnik GmbH, Giessen, Germany
g. Carl Zeiss GmbH, Oberkochen, Germany
h. K. Mußler, Aachen, Germany
i. Sigma Aldrich, Darmstadt, Germany
j. Perkin Elmer, Rodgau, Germany
k. MP Biomedicals Germany GmbH, Eschwege, Germany
l. Roche Diagnostics GmbH, Mannheim Germany
m. Serva Electrophoresis GmbH, Heidelberg, Germany
n. GE Healthcare Europe GmbH, Freiburg, Germany
o. Affinity Biosciences LTD. Cincinnati, OH, US
p. ThermoFisher Scientific GmbH, Dreieich, Germany
q. Merck Millipore, Darmstadt, Germany
r. Bio-Rad Laboratories GmbH, Feldkirchen
s. Heinrich Heine Universität, Düsseldorf, Germany
t. SAS Institute Inc., Cary, North Carolina, USA
u. Graphpad Software Inc., San Diego, California, USA

## Supplementary items

1. Electrophysiology results

## Notes

### Competing Interest Statement

The authors have declared no competing interest.

