## supplementary item 1 for "The effect of ischaemic postconditioning on mucosal integrity and function in equine jejunal ischaemia"

Table 1: Short circuit currents

| | **Control group** | | | **Postconditioning group** | | |
|---|---|---|---|---|---|---|
| | *Pre-ischaemia* | *Ischaemia* | *Reperfusion* | *Pre-ischaemia* | *Ischaemia* | *Reperfusion* |
| *Basal (alanine)* | -0.26 ± 0.2 | 0.32 ± 0.06 | -0.019 ± 0.02 | -0.45 ± 0.6 | 0.28 ± 0.1 | 0.017 ± 0.1 |
| *Alanine* | 3.4 ± 0.8 | 0.17 ± 0.1 | 0.58 ± 0.2 | 3.1 ± 1.1 | 0.68 ± 0.8 | 0.9 ± 1.0 |
| *Forskolin* | 0.56 ± 0.4 | 0.25 ± 0.08 | 0.59 ± 0.2 | 0.61 ± 0.4 | 0.32 ± 0.3 | 0.74 ± 0.6 |
| *Basal (glucose)* | -0.15 ± 0.29 | 0 ± 0.07 | 0.033 ± 0.1 | -0.39 ± 0.4 | 0.032 ± 0.08 | 0.06 ± 0.2 |
| *Glucose* | 2.5 ± 0.6 | 0.11 ± 0.1 | 0.5 ± 0.2 | 2.3 ± 0.8 | 0.52 ± 0.5 | 1.2 ± 0.9 |
| *Forskolin* | 0.52 ± 0.4 | 0.058 ± 0.08 | 0.38 ± 0.2 | 0.61 ± 0.5 | 0.13 ± 0.05 | 0.61 ± 0.4 |

*Table 1 demonstrates the short circuit current in µEq/cm$^2$/h measured in Ussing chambers in intestinal mucosa from horses undergoing ischaemic postconditioning and an untreated control group. The basal values represent the short circuit currents in the alanine and glucose chambers before the addition of these substances (mean ± SD). The measurement after addition of alanine, glucose, or forskolin is expressed in mean change compared to the basal measurement.*

Table 2: Tissue conductance

| | **Control group** | | | **Postconditioning group** | | |
|---|---|---|---|---|---|---|
| | *Pre-ischaemia* | *Ischaemia* | *Reperfusion* | *Pre-ischaemia* | *Ischaemia* | *Reperfusion* |
| *Basal (alanine)* | 16.2 ± 1.7 | 27.6 ± 1.6[a] | 26.3 ± 4.0 | 17.8 ± 4.4 | 22.9 ± 2.6[a] | 24.6 ± 1.5 |
| *Alanine* | 0.2 ± 0.6 | -1.3 ± 0.1[b] | -0.5 ± 0.2 | 0.2 ± 0.4 | -0.1 ± 1[b] | 0.3 ± 1.4 |
| *Forskolin* | 0.3 ± 0.3 | 0.5 ± 0.2 | 0.1 ± 0.3 | 0.4 ± 0.5 | 0.6 ± 0.8 | 0.4 ± 0.7 |
| *Basal (glucose)* | 15.5 ± 0.8 | 21.4 ± 2.9 | 25.1 ± 4.2 | 17.5 ± 3.4 | 23.5 ± 5.0 | 22.6 ± 0.6 |
| *Glucose* | 0.2 ± 0.3 | 0.5 ± 1.0 | -1.1 ± 1.4 | 0.2 ± 0.3 | -0.2 ± 0.4 | 0.0 ± 0.6 |
| *Forskolin* | 0.5 ± 0.3 | 0.6 ± 1.6 | 0.2 ± 0.5 | 0.6 ± 0.4 | 0.1 ± 0.2 | -0.1 ± 0.6 |

*Table 2 demonstrates the tissue conductance in mS/cm$^2$ measured in Ussing chambers in intestinal mucosa from horses undergoing ischaemic postconditioning and an untreated control group. The basal values display the tissue conductance in the alanine and glucose chambers before the addition of these substances (mean ± SD). The values for tissue conductance after addition of alanine, glucose, or forskolin, are expressed in mean change compared to basal measurement.*
